# The superoxide dismutase mimetic GC4419 enhances tumor killing when combined with stereotactic ablative radiation

**DOI:** 10.1101/2020.03.10.984443

**Authors:** Brock J. Sishc, Lianghao Ding, Collin Heer, Debabrata Saha, Douglas R. Spitz, Michael D. Story

## Abstract

The penta-aza macrocyclic manganese compound GC4419 is in phase 3 clinical trials as a modifier of mucositis in H&N cancer treated by radio-chemotherapy based upon its properties as a superoxide dismutase mimetic. In studies to address the potential for tumor radioprotection, a significant anti-tumor effect was identified in tumors generated from the non-small cell lung cancer (NSCLC) cell line H1299, when GC4419 was combined with radiation. This effect was directly related to the size of the radiation dose as demonstrated by greater efficacy in tumor growth delay when biologically equivalent irradiation regimens using a limited number of dose fractions was substantially more effective compared to regimens where the fraction number was higher and dose per fraction decreased. Furthermore, a TCD50 assay using H1299 tumors that tested the combination of GC4419 with radiation revealed a Dose Enhancement Factor of 1.67. Based upon these results the hypothesis that GC4419 was generating cytotoxic levels of hydrogen peroxide during the superoxide dismutation process. Peroxide flux did increase in cells exposed to GC4419 as did the expression of the oxidative stress markers 4-HNE and 3-NT. H1299 cells that overexpressed catalase were then challenged as tumors by the combination of radiation and GC4419 and the tumoricidal effect was nearly eliminated. The enhanced radiation response was not specific to NSCLC as similar findings were observed in human head and neck squamous cell carcinoma and pancreatic ductal adenocarcinoma xenografts. RNA sequencing analysis revealed that GC4419, in addition to generating high levels of hydrogen peroxide in irradiated cells, alters inflammatory and differentiation signaling in the tumor following irradiation. Together, these findings provide abundant evidence that the radioprotector GC4419 has dual functionality and will increase tumor response rates when combined with agents that generate high levels of superoxide like stereotactic ablative body radiotherapy (SAbR). Combining SAbR with GC4419 may be an effective strategy to enhance tumor response in general but may also allow for fully potent radiation doses to tumors that might not necessarily be able to tolerate such doses. The potential for protection of organs at risk may also be exploitable.

## Introduction

Superoxide dismutases (SODs) were first described in 1969 as metalloproteins that dismutated superoxide to hydrogen peroxide and oxygen (*1*). This opened up new avenues of exploration into oxygen toxicity, antioxidant defenses and pathogenic defenses. These enzymes are found in plants and animals within distinct cellular locales including extracellular spaces (SOD3), within the cytosol of cells (SOD1), within the mitochondria (SOD2) and vary by metal group. Overproduction of superoxide or lack of dismutation of superoxide is associated with reperfusion injury (*2–5*), inflammatory processes (*6*), neurodegenerative diseases (*7, 8*), arthritis (*9*), septic shock (*10, 11*), and for a review of the role of superoxides in carcinogenesis see Robbins and Zhao (*12*). Oxidative protein modifications are responsible for the inactivation of proteins involved in sodium channel function, and metabolic and growth regulatory processes (*13, 14*) such as PTEN (*15*), while increased G6PDH activity may mediate superoxide production (*16, 17*). Pre-clinical data from drosophila to mice have identified the protective role of SODs from oxidative damage in a number of settings. See Salvemini (*18*) Robbins, Wang (*14*) and Azadmanesh (*19*) for reviews].

In a review of the role of manganese superoxide dismutase (MnSOD) in cancer, MnSOD was described as diminished in all tumors examined (*20*). Clinical studies have described a link between MnSOD levels and increased cancer incidence in a variety of disease sites and there are numerous studies using knockout or transgenic animals that link MnSOD levels to hepatocellular carcinoma, lymphoma, and others. There are also examples where MnSOD levels are higher such as HNSCC. In this case it has been suggested that higher MnSOD leads to higher levels of H_2_O_2_ that can lead to a variety of oncogenic processes if not detoxified by catalase. See Robbins and Zhao for a review (*12*) The toxicity of hydrogen peroxide is key to this study and will be described below.

SOD mimetics are agents that mimic the activity of naturally occurring SODs. The development and utilization of SOD mimetics is based upon the results of in vitro and *in vivo* studies where cells or animals overexpress naturally occurring SODs or are exposed to recombinant MnSOD. Recombinant MnSOD or MnSOD mimetics has been shown to protect against radiation (*21*) or chemotherapy agent (*22*) damage. See Mupaskar et al for a recent review (*23*). Also important to this study are the studies demonstrating that the addition of MnSOD before application of a carcinogen, while effective at reducing the incidence of carcinogenesis, was even more effective when the MnSOD was applied after application of the carcinogen. While no SOD mimetics are approved for clinical use, a number are in clinical trials. Interestingly, virtually all approaches to the use of SOD mimetics has been as a protective or preventive agent, whether tested in models of ischemia or oxidative damage to the kidney, as examples, or as an anti-carcinogen. However, Hurt et al., (*24*) showed that when SOD2 is overexpressed in pancreatic cancer cells their proliferation slowed; the notion being that this reduction in proliferation was due to redox dependent signaling. This finding too, is relevant to the findings described below.

A recent phase 2b randomized trial demonstrated that vs placebo, the MnSOD mimetic GC4419 substantially reduced oral mucositis in head and neck cancer patients treated with radiation and cisplatin (*25*). (Clinical Trials.gov identifier: NCT02508389). Oral mucositis is a debilitating acute response to radiation that can have considerable impact upon patient outcome. Studies are ongoing in the use of this agent as a defender against lung fibrosis, a radiation-induced late effect, from high dose per fraction radiation protocols like Stereotactive Ablative Radiotherapy (SAbR) as described in Story, et al. (*26*). The application of SAbR protocols is rapidly gaining favor in radiation oncology due to technical innovations from imaging to dose delivery that limit doses to the normal tissue. While initially used in the surgery ineligible lung cancer setting, the use of SAbR has expanded to a number of disease sites because of the outstanding clinical results seen for lung cancer (*27–30*).

Not considered as a clinical strategy is taking advantage of the imbalance in superoxide dismutation seen in cancer cells and the intentional generation of hydrogen peroxide to kill cancer cells. Treatment regimens like SAbR could be expected to generate overwhelming levels of hydrogen peroxide if the superoxide dismutase enzymes are available or can be supplanted with mimetics and the levels of hydrogen peroxide generated overwhelm the already reduced levels of catalase often seen in tumors. That strategy was employed in this study where the SOD mimetic was tested as an anti-cancer agent when combined with high dose per fraction radiation schemes in a series of tumor types. Indeed, the anti-tumor efficacy was correlated with increasing radiation dose and was abrogated by the upregulation of catalase in the tumors designed to overexpress catalase using an inducible promoter system. Moreover, a Tumor Control Dose (50%) assay determined that there was a dose enhancement factor was 1.67 which is generally more than that seen with other agents used in combination with radiation: gemcitabine 1.4-1.54 (*31, 32*); C225 1.8 (*33*); cisplatin 1.3-1.4 (*34*); nimorazole 1.3 (*35*), as examples. The use of GC4419 in combination with SAbR represents the promise of offering fully potent treatments to large tumors, proximal tumors of the lung or other tumors near organs at risk and radioresistant tumors. A clinical trial combining GC4419 and SAbR for inoperable pancreatic cancer is ongoing (Clinical Trials.gov identifier: NCT03340974) while a trial in lung cancer will commence shortly. Lastly, what is quite novel is the potential for both radioprotection and an enhanced tumor response in the context of SAbR. Pre-clinical studies of both radioprotection of normal tissue, enhanced tumor response and metastatic impact in the same model system is ongoing.

## Results

### GC4419 does not protect human non-small cell lung cancer (NSCLC) cell lines or tumors from radiation

Given the reduction of acute radiation-induced mucositis using the hamster cheek pouch model (*36*) and a precursor compound to GC4419, it was important to test tumor cells in vitro and in vivo for the potential for tumor radioprotection. The chemical structure of GC4419 is depicted in Figure 1A. First, *in vitro* clonogenic survival assays were conducted using four NSCLC cell lines. At supra-physiological concentrations of GC4419 (the maximum concentration in tissues peaks at 10 μM) NSCLC cells are not protected, and a slight sensitization, although not statistically significant, is observed (Figure 1). This observation was consistent across all cell lines regardless of their genetic background and is described in supplemental Table 1. Second, tumor growth delay experiments (TGD) were conducted where a single dose of GC4419 was administered 30 minutes prior to the irradiation of H1299 cells grown as tumors in the leg of athymic nu/nu mice.

**Figure 1:**
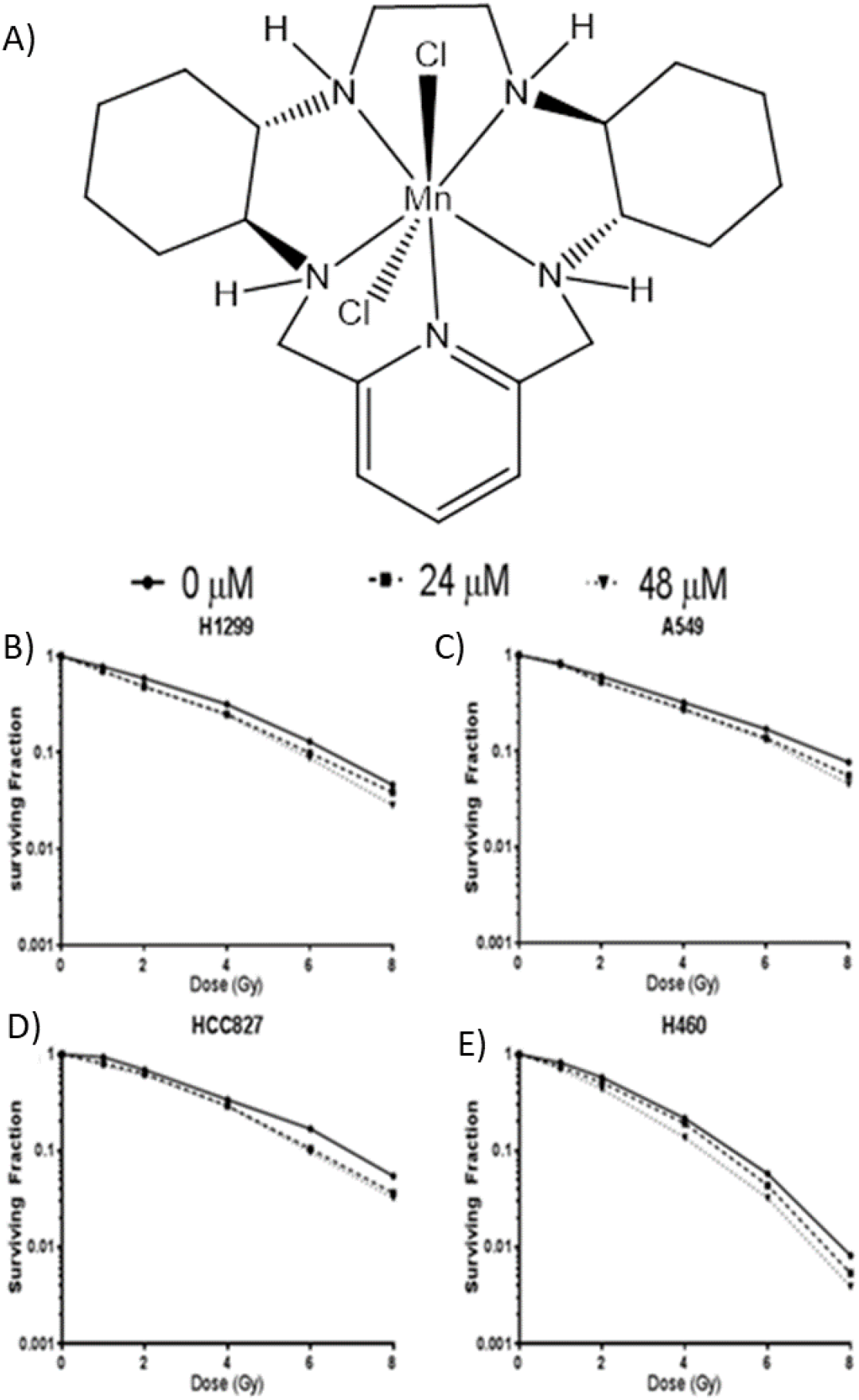
Pre-treatment of NSCLC cell lines with GC4419 does not protect against radiation induced damage. A) Chemical structure of GC4419. NSCLC tumor cell lines B) H1299, C) A549, D) HCC827, and E) H460 were treated with supraphysiological 24 and 48 μM concentrations of GC4419 30 minutes prior to irradiation with 0-8 Gy of γ-rays. Dose response curves are shown to demonstrate that GC4419 not only does not protect these cell lines from radiation, but it also results in slight sensitization of the cells to ionizing radiation.

Synergistic enhancement of the response of the tumors to a single fraction of 18 Gy was observed with a dose enhancement factor (DEF) of 1.6 (Figure 2 A-B). And, to determine whether drug half-life might play a role in tumor radioresponse, a second experiment was conducted where GC4419 was delivered once 30 minutes prior to irradiation and an additional daily dose was delivered on each of the four consecutive days (5 daily doses of GC4419 total) (Figure 3 C-D). Not only did GC4419 synergistically enhance the response of tumors to radiation, the DEF could not be calculated because the result was tumor cure. This finding was repeated using two additional human tumor xenografts, A549 and HCC827 (Figure 3 E-L). Data is depicted as tumor average tumor volume as well as the volume of individual animals as a function of time. Furthermore, GC4419 demonstrated mildly enhanced tumor growth delay (TGD) as a single agent.

**Figure 2:**
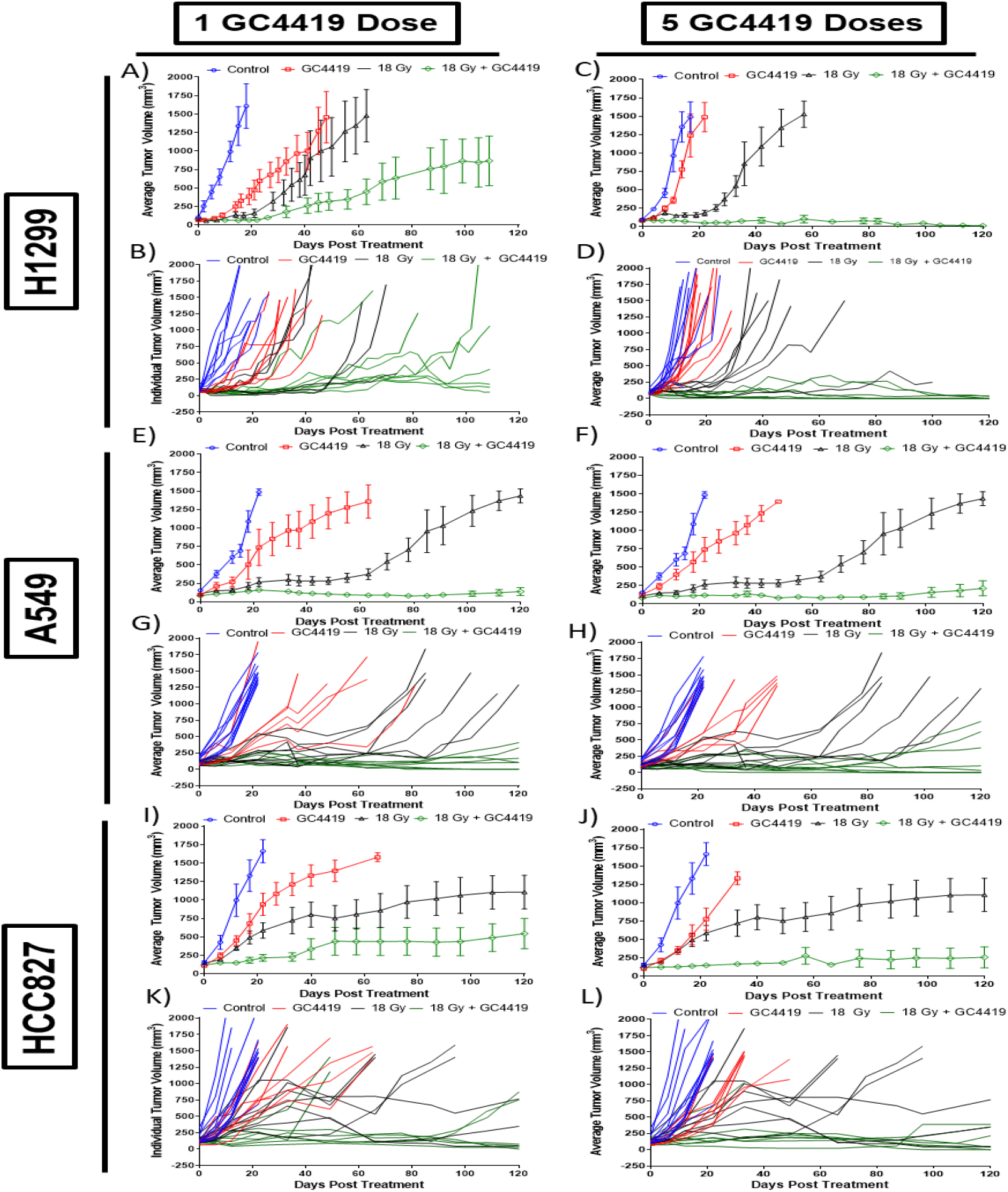
Tumor growth delay experiments in H1299, A549, and HCC827 ectopic xenografts combining IR and GC4419. Averaged (top panels) and individual (bottom panels) tumor volumes comparing the effects of GC4419 alone, and combined with a single dose of 18 Gy. H1299 (top panel), A549 (middle panel), and HCC827 (bottom panel) were all evaluated with a single dose of GC4419 30-60 minutes prior to irradiation (left column) and GC4419 delivered on an additional 4 days following irradiation (right column).

**Figure 3:**
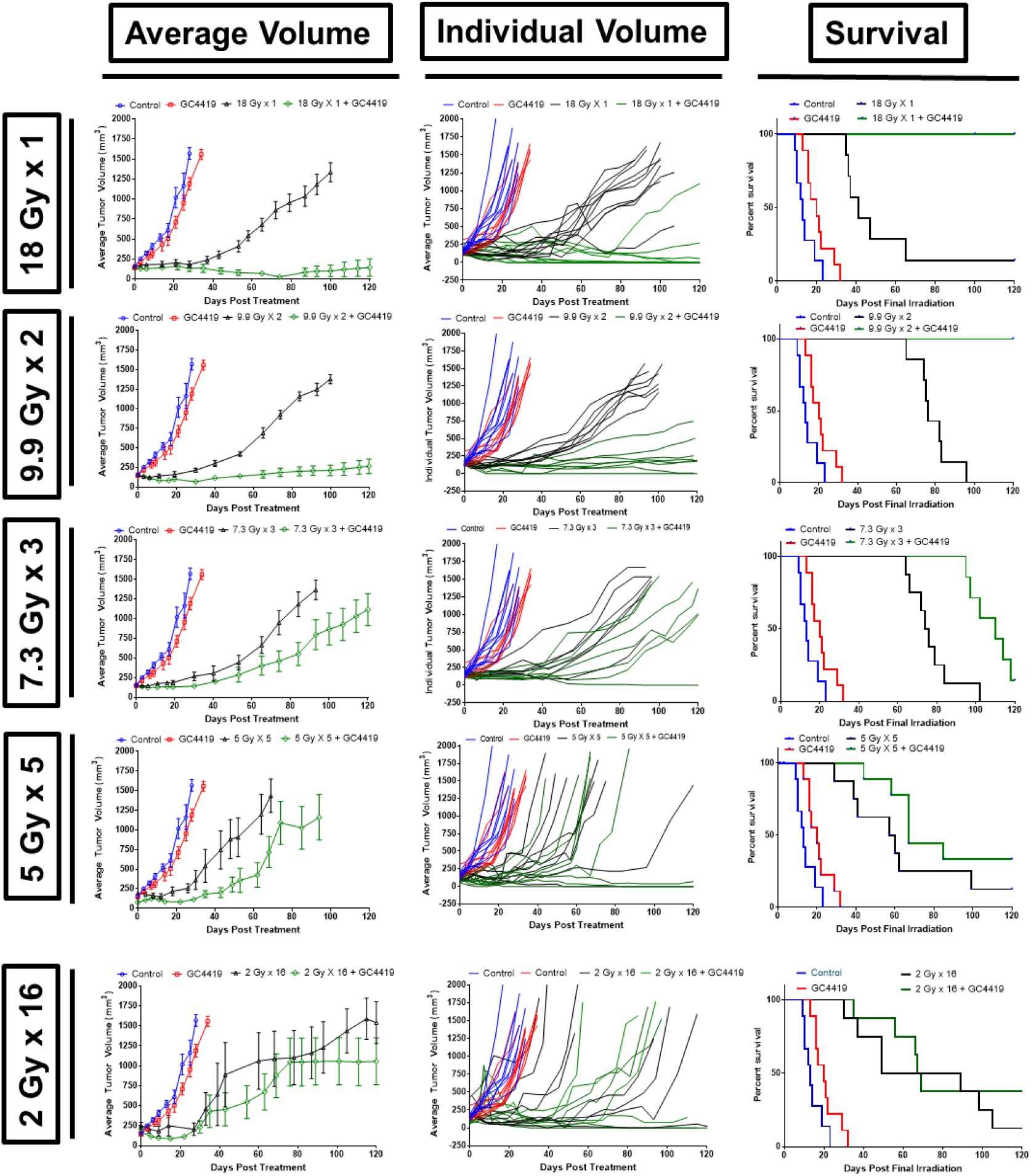
GC4419 is more effective at enhancing the radiation response at high doses per fraction. H1299 xenorafts were generated to determine whether GC4419 was more effective in the high dose per fraction setting. Schedules of 18 Gy × 1 fxn, 9.9 Gy × 2 fxn, 7.3 Gy × 3 fxn, 5 Gy × 5 fxn, and 2 Gy × 16 fxn were compared. Average tumor volumes (left column), individual tumor volumes (middle column), and Kaplan Meier analysis using the IACUC threshold of 1000 mm^3^ tumor volume as a proxy for survival.

### The enhanced radiation response with GC4419 is more pronounced at high dose per fraction

The dose of 18 Gy chosen for H1299 TGD experiment was chosen so as to reflect any potential for tumor radioprotection or radiosensitization. However, the overwhelming anti-tumor effect seen in Figure 2 suggested the potential for an additional mechanism of tumor cell killing that relied on high dose per fraction radiation exposures. To test this notion, biologically equivalent fractionation schedules equivalent to the 18 Gy × 1 fraction (Figure 3 A-C), consisting of 9.9 Gy × 2 fractions (Figure 3 D-F), 7.3 Gy × 3 (Figure 3 G-I) fractions, 5 Gy × 5 fractions (Figure 3 J-L), and 2 Gy × 16 (Figure 3 M-O) fractions delivered on consecutive days, were selected. These fractionation schemes yielded equivalent tumor growth delay curves. Average tumor volume, individual tumor volumes, and Kaplan Meier survival curves for these five fractionation schedules using the 1000 mm^3^ IACUC tumor size threshold as a proxy for survival are displayed. Interestingly, GC4419 is more effective at enhancing the radiation response when used with higher doses per fraction, with a threshold between 7.3 and 9.9 Gy. While not significant statistically, there is a slight enhancement of the radiation response even in the 5 Gy × 5 fractions group. Additionally, GC4419 does not protect tumors exposed to 2 Gy × 16 fractions.

### GC4419 increases the rate of tumor cure with a dose enhancement factor of 1.67

Tumor growth delay is not a true measure of tumor cure, as any single surviving clonogen can effectively repopulate the tumor resulting in locoregional failure, in addition to the potential for acquired therapeutic resistance. Therefore, to determine the potential for GC4419 to enhance tumor cure rate, tumor cure dose to achieve 50% tumor kill (TCD_50_) assays were performed in ectopic H1299 xenografts with radiation alone and radiation in combination with GC4419 at a 24 mg/kg dosage (Figure 4). GC4419 shifted the TCD_50_ dose from 25 Gy with radiation alone to 15 Gy, resulting in a dose enhancement factor (DEF) of 1.67. Furthermore, GC4419 also altered the slope of the TCD_50_ curve, indicating that not only does it enhance the efficacy of radiation, but it does so by altering the biological response of the tumor to radiation, altering the γ-function.

**Figure 4:**
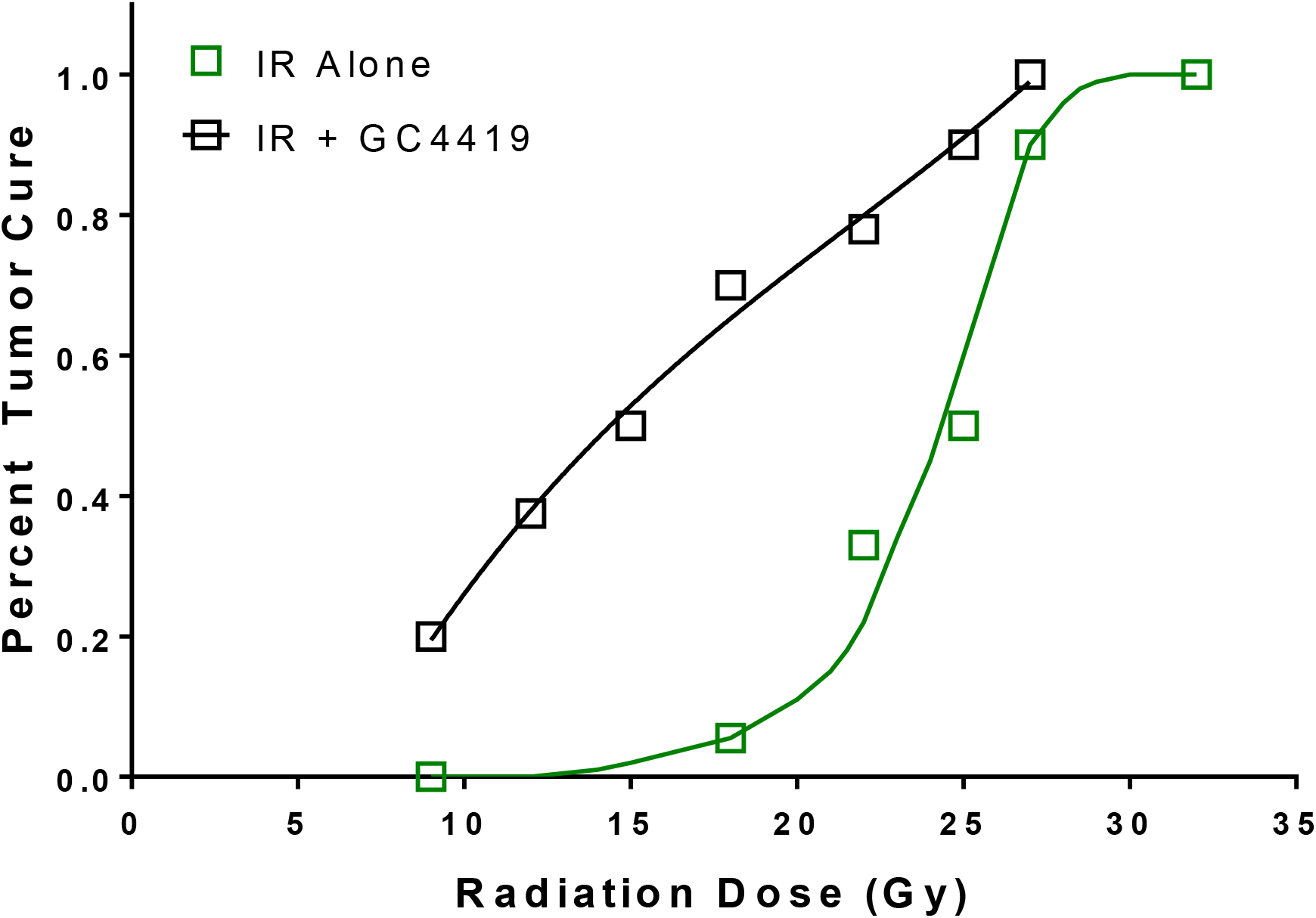
GC4419 shifts the TCD_50_ dose of H1299 xenografts to radiation exposure with a dose enhancement factor of 1.67. H1299 tumors were treated with curative doses of radiation with or without GC4419 and followed for 120 days post radiation exposure.

### Overexpression of catalase in H1299 xenografts diminishes the enhanced radiation response seen with GC4419

Given the rate at which GC4419 dismutates superoxide to hydrogen peroxide, it seemed reasonable to propose that increasing levels of hydrogen peroxide would be produced with increasing doses of radiation. In fact, without radiation the addition of GC4419 to H1299 cells in culture did increase the flux of hydrogen peroxide (Figure 5A). Therefore, scavenging elevated levels of hydrogen peroxide produced by GC4419 superoxide dismutation would blunt the enhanced radiation response observed with GC4419. To test this hypothesis, a H1299 tumor cell line that overexpresses catalase on a doxycycline inducible promoter was utilized (H1299-CAT). Figure 5 demonstrates that H1299-CAT tumors have elevated catalase activity when doxycycline is delivered to tumor bearing animals *ad libitum* in drinking water (2.5 mg/mL) (Figure 5B). TGD experiments demonstrate that without doxycycline in the drinking water, and therefore no overexpression of catalase, H1299-CAT tumor growth delay had a similar response to the H1299 parent line in that GC4419 had a mild anti-tumor effect alone, and synergistically enhanced the effects of ionizing radiation (Figure 5 C-D). However, when doxycycline was added to the drinking water and catalase was overexpressed, the enhanced radiation response observed with GC4419 was blunted (Figure 5 E-F). As further evidence for the reduction of oxidative stress, the expression of 4HNE and 3-NT, both markers of oxidative stress, was determined in H1299-CAT tumors treated with or without GC4419 while overexpressing catalase. The activation of catalase reduced both the expression of 4HNE and 3-NT, suggesting that catalase overexpression is repressing oxidative stress (Supplemental figure 1A-B).

**Figure 5:**
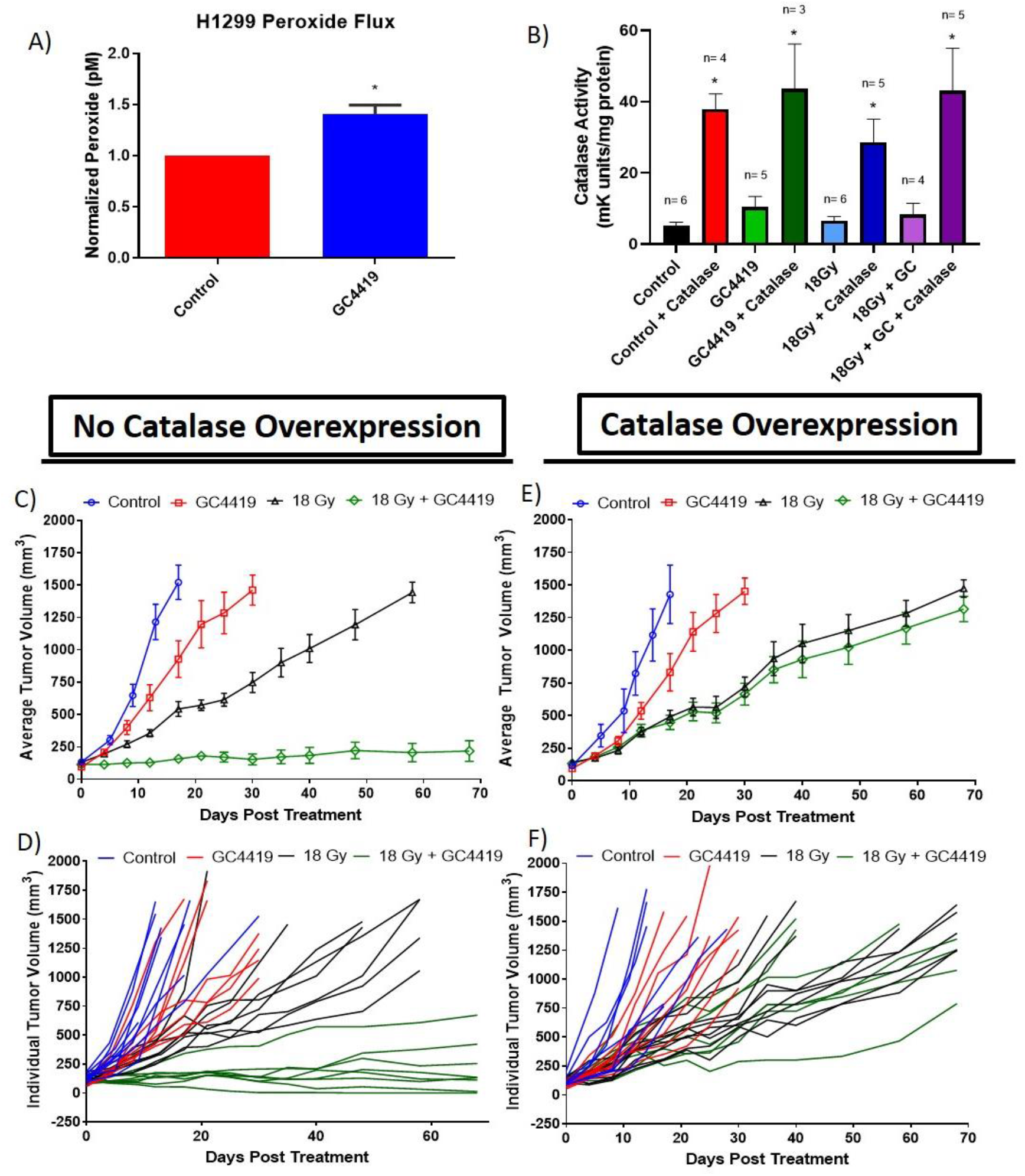
Overexpression of catalase eliminates the ability of GC4419 to enhance the radiation response. A) GC4419 increases hydrogen peroxide flux when added to H1299 cells in culture. H1299-CAT tumors were modified to overexpress catalase on a doxycycline inducible promoter. B) The induction of catalase activity in H1299-CAT cells by treatment with doxycycline *in vitro*.T GD of animals exposed to a combination of 18 Gy and GC4419 without (left) or with doxycycline (right) in the water (n = 8 animals per group). Doxycycline induced overexpression of catalase eliminated the enhanced radiation response observed with GC4419 on both the average (C/E) and individual basis (D/F)).

### The enhanced response of tumors to radiation with GC4419 is not specific to NSCLC

Many tumor types have decreased levels of catalase expression and activity (*20*), a hydrogen peroxide based cell death mechanism presented the possibility that the enhanced response of tumors to radiation treatment was not specific to NSCLC. Therefore, additional xenografts were generated using four additional human tumor lines. The radioresistent SqCC/Y1 HNSCC cell line was selected because of the results of the aforementioned phase IIB clinical trial where GC4419 demonstrated radioprotective efficacy in patients receiving treatment for head and neck cancer (Clinical Trials.gov identifier: NCT02508389). The PDAC cell lines Panc 02.03, PANC-1, and SW1990 tumor lines were selected due to the initiation of a Phase I/II clinical trial examining a potential role for GC4419 to enable dose escalation in patients receiving SAbR for the treatment of pancreatic cancer (Clinical Trials.gov identifier: NCT03340974). TGD studies were conducted using a single dose of 12 Gy. Again, GC4419 treatment resulted in mild (albeit to varying degrees) anti-tumor activity as a single agent and enhanced the response of SqCC/Y1 (Figure 6 A-B), Panc 02.03 (Figure 6 C-D), SW1990 (Figure 6 E-F), and PANC-1 (Figure 6 G-H) tumors to radiation.

**Figure 6:**
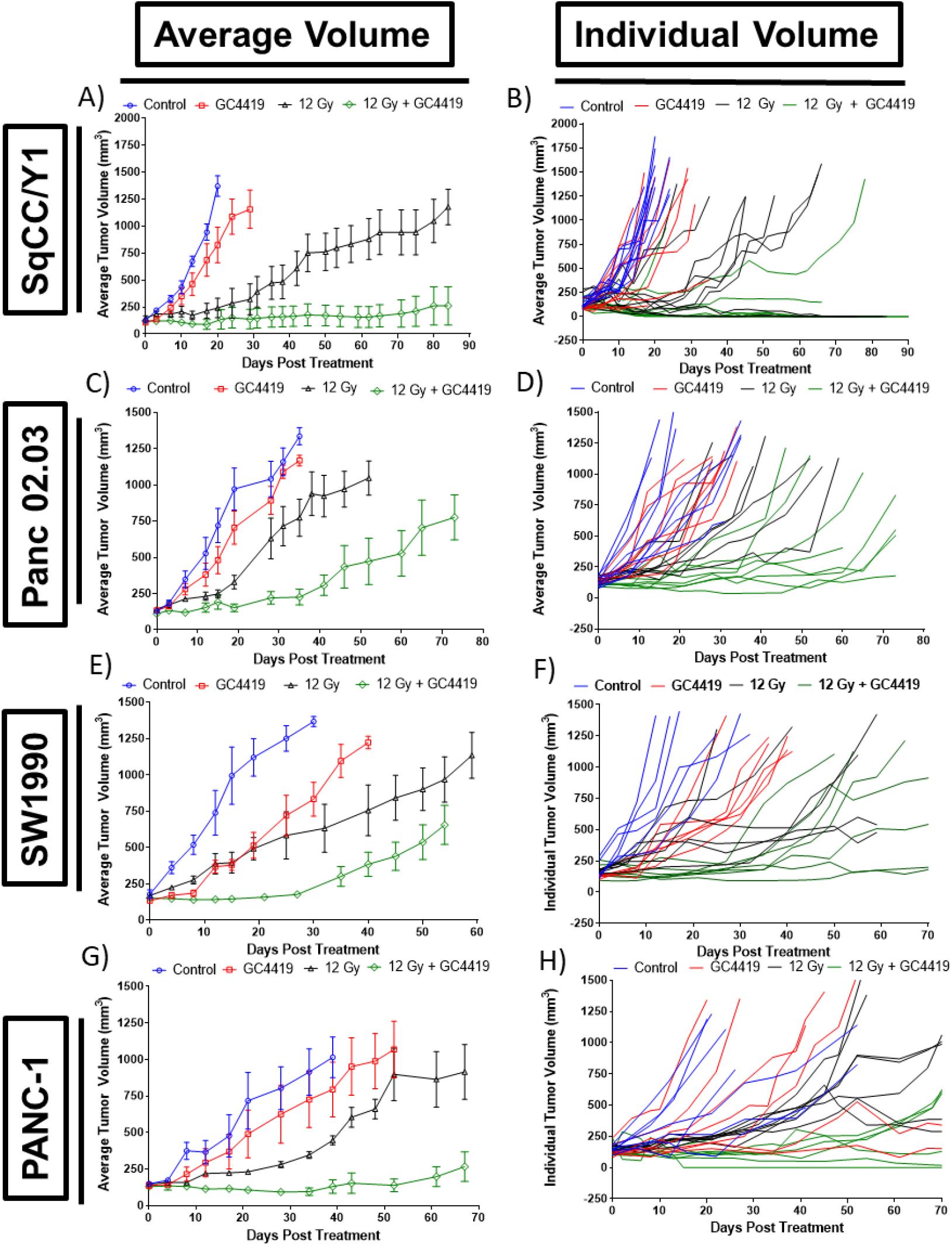
Enhancement of the response of tumors to radiation is not specific to NSCLC xenografts. GC4419 was tested alone and in combination with a single dose of 12 Gy X-rays using cell lines SqCC/Y1 (HNSCC), Panc 02.03, SW1990, and PANC-1 (all PDAC). Average tumor volumes (left column) and individual tumor volumes (right column) are displayed.

### Transcriptome profiling of H1299 xenografts by treatment group

Total RNA sequencing was performed on H1299 tumors treated with an 18 Gy radiation dose with or without GC4419 given 30 minutes prior to radiation and daily for an additional 4 days. RNA was extracted from tumors on days 0, 1, 3, and 7 following the initiation of treatment. Unsupervised gene enrichment analysis identified 7 pathways that were significantly changed when comparing the irradiated tumors with or without the addition of GC4419. Pathways identified as up-regulated in the radiation plus GC4419 combination group included the myogenesis pathway, TNF☐ signaling via NFKB, EMT signaling, inflammatory response, androgen response, hypoxia signaling and apoptosis signaling. Interestingly, one pathway, hedgehog signaling, was significantly down-regulated in the combination group (Supplemental Table 2). Normalized expression values from the leading edge subset of genes in the above pathways were plotted as heatmaps to show detailed changes over the time course (Figure S2-3). Expression values calculated from the heatmaps show pathway changes differed across time. Hedgehog signaling, was initially reduced in the radiation plus GC4419 group compared to radiation only, with signaling for both radiation and radiation plus GC4419 increasing with time. Inflammatory signaling appeared to be higher earlier in the radiation plus GC4419 cohort. Most pathways by day 7 saw little difference in expression between the two groups which may reflect drug availability.

Supervised expression analysis identified 388 genes, including 62 long non-coding RNA that exhibited different expression profiles based upon treatment group status and time post irradiation by Principal Component Analysis (PCA) seen in Figure 7A. PCA demonstrated that, while the gene expression patterns of tumors sampled earlier in the treatment schedule were not well separated from the untreated control tumors, by day 7 the different groups could be found within their own three-dimensional space. There was, however, significant overlap between the radiation and radiation plus GC4419 group at day 7. Pathway analysis was performed using the 388 genes from the PCA. Those results identified significantly altered canonical pathways including metabolic and signaling pathways, dependent on different treatment groups over the time course (Figure 7B).

**Figure 7:**
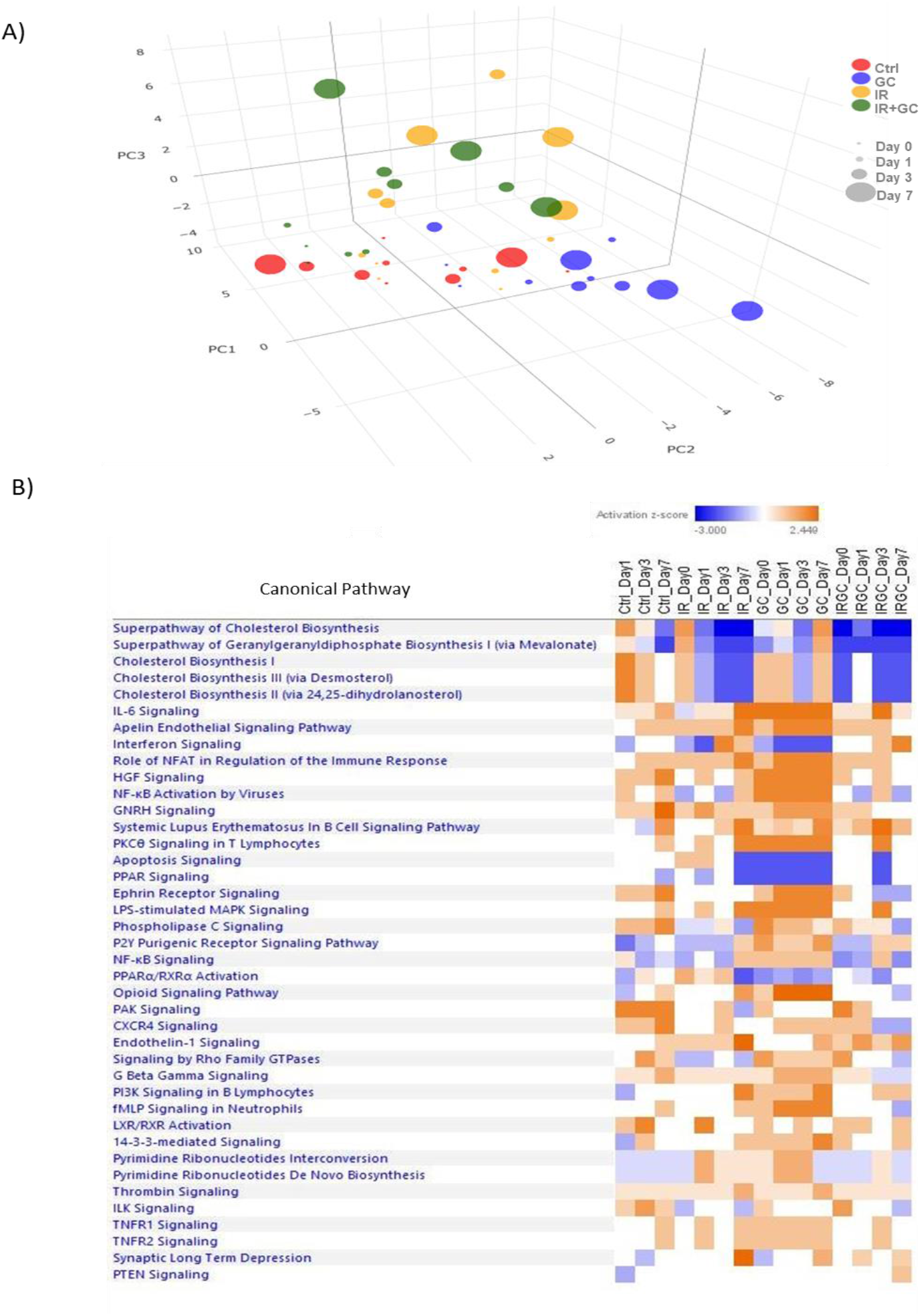
Identification of a 388 gene signature that spatially separates treatment groups. A) Principle component analysis showing the separation of the four treatment groups (control, GC4419, 18 Gy, and 18 Gy + GC4419). Colors represent treatment group while the size of each individual dot represents the time after treatment that the tumor tissue was collected. B) Canonical pathways identified as altered when comparing treatment groups with the described 388 gene signature.

## Discussion

Not only does GC4419 not protect NSCLC cells and tumors from IR exposures, GC4419 significantly enhances the response of NSCLC tumors to irradiation in vivo when radiation is delivered at high dose per fraction. This enhanced response, as measured by TGD is independent of genetic background and tumor type. Furthermore, this effect leads to a decreased IR dose threshold necessary to eliminate all the clonogens within the tumor, effectively resulting in higher tumor cure rates at lower doses.

Ionizing radiation exposure results in three waves of oxidative stress generation; immediate; early; and a chronic phase that contributes to long lasting oxidative stress. The immediate response phase results from the radiolysis of water producing superoxide and other reactive species, and is short lived. The early phase results from the upregulation of NADPH oxidase activity which increases the levels of superoxide from 1-24 hours post IR exposure (*37, 38*). The chronic phase, likely resulting from mutations in mitochondrial electron transport chain genes, alters the polarization of mitochondria leading to increased levels of superoxide that can even be passed on to daughter cells (*39*). These elevated levels of biologically produced superoxide are proportional to IR dose, indicating that a threshold may be required to induce this phenomenon (*40*).

The upregulation of SODs can spare tumor cells from these chronic effects, however, the unique chemistry of GC4419 which is both rapid and a true catalyst, combined with the downregulation of catalase in most tumors led to the hypothesis that the increased levels of superoxide resulting from high dose per fraction irradiation are in fact being immediately converted to toxic levels of hydrogen peroxide. Indeed, numerous studies have demonstrated that increasing expression of SOD in tumor lines have anti-tumor properties (*41–47*), a finding enhanced by inhibiting catalase (*47*). Overexpression of catalase, the primary but not only hydrogen peroxide detoxifying enzyme, effectively eliminated the enhanced radiation response when tumors were treated with radiation and GC4419. This finding is consistent with recent reports that suggest that elevated levels of hydrogen peroxide either produced by SOD mimetics, or by altering metabolic processes to produce more hydrogen peroxide, is an effective strategy in cancer therapy (*48*).

Given the reduction in acute mucositis seen in the hamster cheek pouch model and in the human head and neck clinical trial (*25, 36*) and the anti-tumor response seen here when high dose per fraction radiation was used, there is reason to believe that there is a dual functionality for GC4419 as both a radioprotector of acute radiation-induced normal tissue toxicity and as an enhancer of the anti-tumor radiation response. Normal tissues, with higher levels of radical oxygen scavenging capacity are likely better able to lessen the free radical damage generated by radiation per se as well as the hydrogen peroxide toxicity produced by the dismutation of superoxide by GC4419. In the tumor however, where the redox capacity of cells is reduced, as noted by reduced levels of the catalase enzyme seen in most tumor cells (*20*), GC4419 produces excess hydrogen peroxide through the dismutation of radiation-induced superoxide (immediate) as well as biological superoxide production (early and chronic) that is upregulated from high dose per fraction radiation exposure. This likely explains the additional cell killing seen with additional doses of GC4419 post-radiation. Interestingly, adding an additional 9 doses of GC4419 instead of 4 doses did little to increase cell killing (data not shown).

Taking clinical advantage of this dual functionality for GC4419 is likely dependent upon the intent and thus strategy of application. For example, in the context as both a modifier of acute adverse events, perhaps where high dose per fraction radiotherapy is not prudent because of the risk for adverse events, GC4419 has a role that may actually allow for dose escalation to a tumor using conventional radiation doses. In the context of enhancing the anti-tumor effects of high dose per fraction radiotherapy, GC4419 has the potential to perform as both as an anti-tumor agent, but also as a modifier of potential radiation-induced adverse normal tissue events. Finally, as the data within show, the mechanism of action is likely not specific to NSCLC and clinical trials should be conducted examining the potential for GC4419 to enhance radiotherapy at other tumor sites where SAbR is an effective treatment strategy.

## Materials and Methods

### Cell Lines and Cell Culture

Cell lines H1299, A549, H460, HCC827, PANC-1, Panc 02.03, and SW1990 were purchased from the American Type Cell Culture repository. The SqCC/Y1 cell line was a kind gift from Dr. Katherine Mason, MD Anderson Cancer Center. Human lung adenocarcinoma cell lines H1299, A549, H460 and HCC827 were cultured in DME/F12 basal media (Sigma Aldrich) supplemented with 10% Fetal Plus fetal bovine serum (FBS) (Atlas Biologicals) and L-Glutamine (Sigma Aldrich). Human pancreatic ductal adenocarcinoma (PDAC) lines PANC-1 and SW1990 and human squamous cell carcinoma of the head and neck (HNSCC) line SqCC/Y1 were cultured in Dulbecco’s Modified Essential Medium with high glucose (Sigma Aldrich) supplemented 10% FBS and L-Glutamine. PDAC cell line Panc 02.03 was cultured in RPMI-1640 basal media (Sigma Aldrich) supplemented with 15% FBS, L-Glutamine, and 10 Units/mL human recombinant insulin (Gibco). All cell culture was conducted in a class two, certified biological safety cabinet and cells were grown at 37^oC^ at 5% CO_2_ in a humidified incubator.

### Tumor Growth Delay and TCD_50_ Studies

Animals were purchased from Charles River and maintained and handled under protocols approved by the Institutional Animal Care and Use Committee (IACUC) at the University of Texas Southwestern Medical Center. For experiments using human tumor xenograft lines, 6 week old female athymic nu/nu mice were inoculated with 5 × 10^6^ human tumor cells (H1299, A549, HCC827, SqCC/Y1, Panc 02.03, SW1990, PANC-1, and SqCC/Y1) suspended in serum free culture medium subcutaneously to the right hip. Tumor volume was measured using calipers 2-3 times weekly. Tumor treatment was initiated when tumor volume ranged between 100-150 mm^3^ and volume was tracked until euthanasia or 120 days post treatment, at which point if no mass was present, the tumor was considered “cured.”

### Irradiations

Irradiations for *in vitro* cell culture experiments were performed using a Mark 1 sealed ^137^Cs source irradiator (J.L. Shepherd and Associates). Animal irradiations were performed using a X-Rad 320 irradiator (Precision X-ray) running at 250 kVp and 15 mA. Tumor irradiations were performed while animals were under anesthesia and radiation was targeted to focally irradiate the tumor utilizing a 10 mm collimated beam to spare at risk normal structures. For studies using fractionated exposures, fractions were delivered on consecutive days.

### Drug Handling and Delivery

GC4419 was provided by Galera Therapeutics and was solubilized to a concentration of 24 mM in bicarbonate buffered saline solution at a pH of 7.4. GC4419 was delivered as a single injection intraperotineally as a 24 mg/Kg dose 30-60 minutes prior to irradiation. Doxycycline hydrochloride was purchased from VWR and solubilized to a concentration of 2.5 mg/mL for animal experiments in distilled deionized water with 1% sucrose.

### Clonogenic Survival Assays

Cells were plated into 60 mm^2^ dishes and allowed to adhere for 6 hours. Following attachment, cells were irradiated with γ-ray doses of 2, 4, 6, and 8 Gy. Cells were returned to incubation and allowed to divide for 10 population doublings (7-9 days depending on cell line). Plates were then stained with 0.5% crystal violet solution in dissolved in 20% methanol. Colonies were counted using a dissecting microscope and surviving fraction was determined based on the number of colonies containing 50 or more cells. Results indicate the compilation of three individual biological replicates.

### Generation of Catalase Overexpressing H1299 Cells

H1299 parent cells were transduced using lentivirus to express the CAT gene on a doxycycline inducible CMV promoter. Following viral transduction, cells were selected in puromyocin at a concentration on 2 μM until colonies had formed. Cell lines were then generated from clonal isolates and tested for catalase activity. Validation of the upregulation of catalase was confirmed using a colorimetric assay (Santa Cruz Biosciences).

### Collection of Tumors for total RNA sequencing analysis and Isolation of Tumor Total RNA

Tumors from treated animals were collected on days 0, 1, 3, and 7 following irradiation or initiation of GC4419 treatment. H1299 tumor tissue was collected and 25 mg of tumor was homogenized in Qiazol lysis reagent (Qiagen) and samples were immediately flash frozen in liquid nitrogen and stored at −80^OC^. Once all samples had been collected, a miRNeasy Mini Kit (Qiagen) was utilized to isolate total RNA. Once isolated, RNA concentration was determined using a Nanodrop 2000 Spectrophotometer (Thermo Scientific) and quality was assessed using an Experion Electrophoresis system (Bio-Rad).

### Transcriptome Analysis of Tumor RNA

Sequencing library was prepared using the Illumina (San Diego, CA) Truseq Stranded Total RNA Prep Kit. Single end next generation sequencing was performed by DNAlink (San Diego, CA) using an Illumina Nextseq500 sequencer with an average coverage of 48.6M. Sequencing reads were trimmed to remove adaptor sequences and were aligned with human genome GRCh38 using STAR with the 2-pass option. The aligned reads were quantified using the RSEM program. Gene counts were imported into DESeq2 using tximport. Data processing was performed using the Stampede2 supercomputer at the Texas Advanced Computing Center (TACC). Genes with very few reads were removed by filterByExpr function within edgeR package.

Gene Set Enrichment Analysis was performed using GSEA 4.0.1. Genes from the whole transcriptome profiles were ranked based on p-values derived from Wald’s test comparing the radiation alone group and the radiation plus GC4419 group. Top GSEA Hallmark pathways were identified by selecting q-values < 0.4.

Supervised gene lists that differentiated experimental cohorts in this study were obtained by the time series analysis using DESeq2 with a fdr < 0.2. Gene counts were transformed before PCA analysis using the variance stabilizing transformation (vst) function in DESeq2 package. Pathway analysis of the supervised gene list was performed using Ingenuity Pathway Analysis software (Qiagen Inc.). Top canonical pathways were selected by applying a FDR < 0.05 and z-score > 1 in Fisher’s Exact Test. Normalized counts for each gene were centered and converted to z-scores to generate heatmaps.

### Statistical Analyses

For all animal experiments, the number of animals per group necessary to generate results was calculated to have an α of 0.05 and 80% power. For experiments where TGD is the endpoint, the log rank test was utilized to determine significance.

---

**Supplemental Table 1:** Genetic background of common mutations in NSCLC tumors.

| Cell Lines | EGFR | LKB1<br>STK11 | Kras | H-Ras | N-Ras | P53 | PTEN | EML4-ALK | SF2 |
| --- | --- | --- | --- | --- | --- | --- | --- | --- | --- |
| A549 | WT | Null | Mut | WT | WT | WT | WT | No | 0.74 |
| H460 | WT* | Null | Mut | WT | WT | WT | WT | Yes | 0.54 |
| H1299 | WT | Null | WT | WT | Mut | Null | WT | No | 0.75 |
| H1993 | WT |  | WT | ? | WT | Null | WT | No | 0.38 |
| H2228 | WT |  | WT | ? | WT | Null | WT | Yes | 0.20 |
| H3122 | WT |  | WT | ? | WT | Null | WT | Yes | 0.56 |
| H1975 | Mut<sup>1</sup> |  | WT | WT | WT | Null | WT | No | 0.38 |
| HCC827 | Mut<sup>2</sup> |  | WT | WT | WT | Null | WT | No | 0.62 |
| HCC4017<sup>4</sup> |  | Null | 12TGT |  | WT | Null | WT |  | 0.32 |

**Supplemental Table 2:** Top Hallmark Pathways Changed Between 18 Gy and 18 Gy + GC4419 Cohorts.

| Pathway | ES | NES | p-value | q-value | FWER p-value | Up or Down in IR+GC |
| --- | --- | --- | --- | --- | --- | --- |
| HALLMARK_MYOGENESIS | 0.43 | 1.43 | 0.001 | 0.329 | 0.298 | UP |
| HALLMARK_TNFA_SIGNALING_VIA_NFKB | 0.4 | 1.37 | 0.001 | 0.349 | 0.534 | UP |
| HALLMARK_EPITHELIAL_MESENCHYMAL_TRANSITION | 0.39 | 1.34 | 0.003 | 0.349 | 0.68 | UP |
| HALLMARK_INFLAMMATORY_RESPONSE | 0.4 | 1.34 | 0.015 | 0.265 | 0.685 | UP |
| HALLMARK_ANDROGEN_RESPONSE | 0.41 | 1.31 | 0.042 | 0.272 | 0.769 | UP |
| HALLMARK_HYPOXIA | 0.36 | 1.22 | 0.045 | 0.386 | 0.984 | UP |
| HALLMARK_APOPTOSIS | 0.36 | 1.21 | 0.056 | 0.387 | 0.99 | UP |
| HALLMARK_HEDGEHOG_SIGNALING | 0.61 | 1.73 | 0 | 0.008 | 0.008 | DOWN |

**Supplemental Figure 1:**
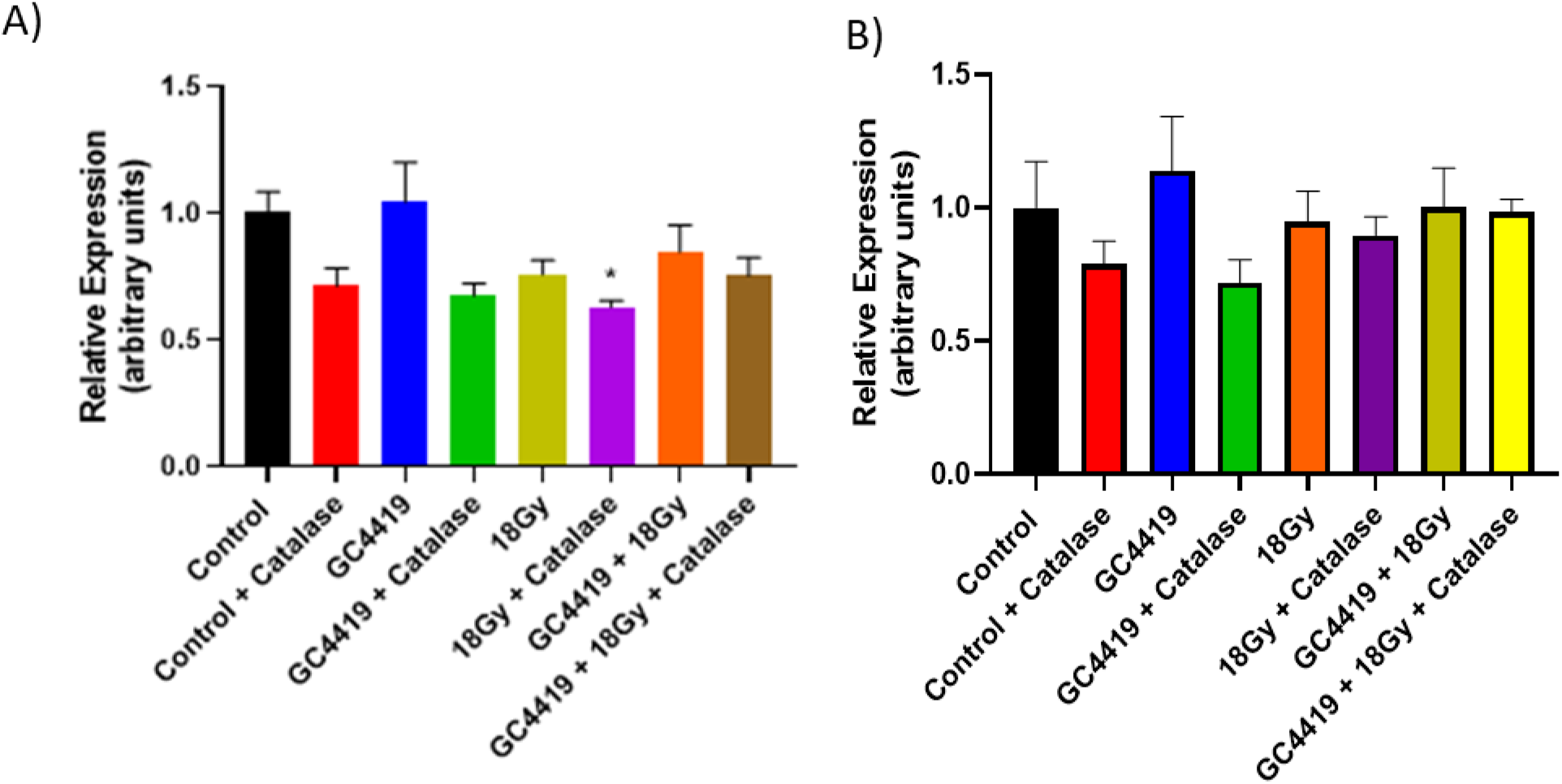
Expression of oxidative stress markers 4-HNE and 3-NT in H1299 and H1299-CAT tumors. Expression of 4-HNE (A) and 3-NT (B) in tumors overexpressing catalase or not at 3 days post treatment start.

**Supplemental Figure 2:**
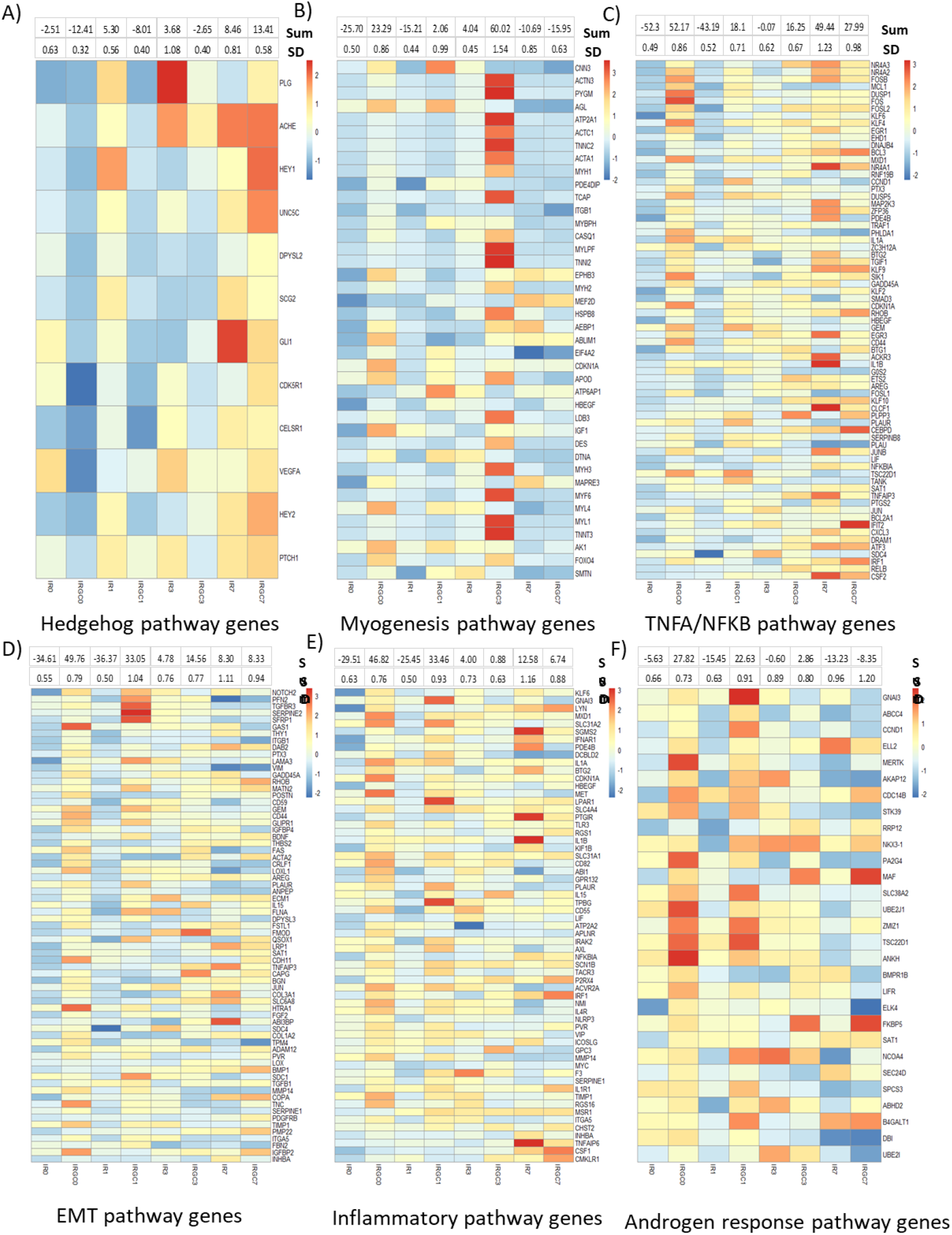
Heatmaps from supervised cluster analysis depicting differentially regulated pathways. Heatmaps representing the sum of Z-scores for each pathway by treatment group for hedgehog signaling (A), myogenesis (B), TNF-α/NFκB signaling (C), EMT (D), inflammatory signaling (E), and androgen response (F).

**Supplemental Figure 3:**
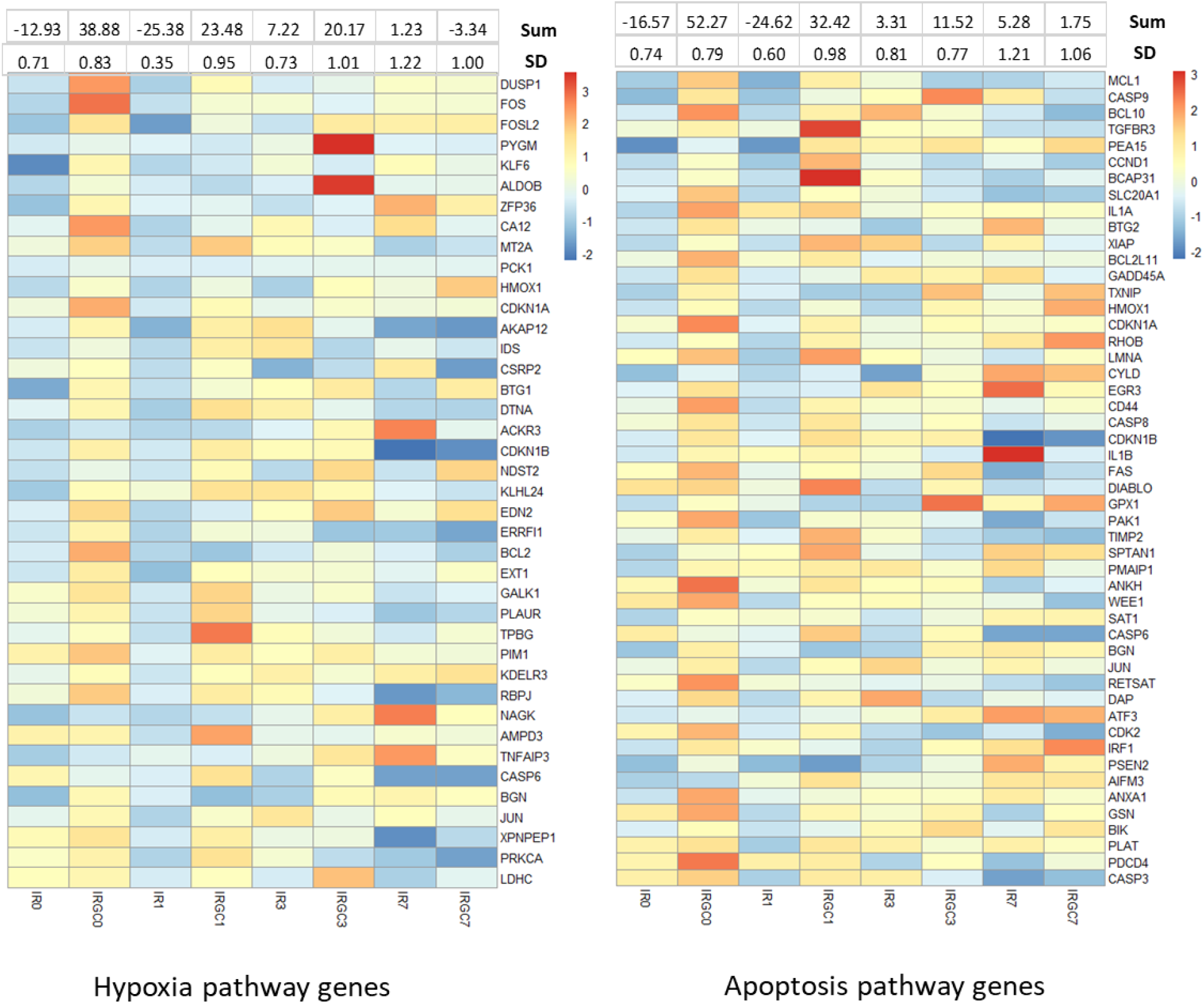
Heatmaps from supervised cluster analysis depicting differentially regulated pathways. Heatmaps representing the sum of Z-scores for each pathway by treatment group for hypoxia signaling (A) and apoptosis (B).

**Supplemental Figure 4:**
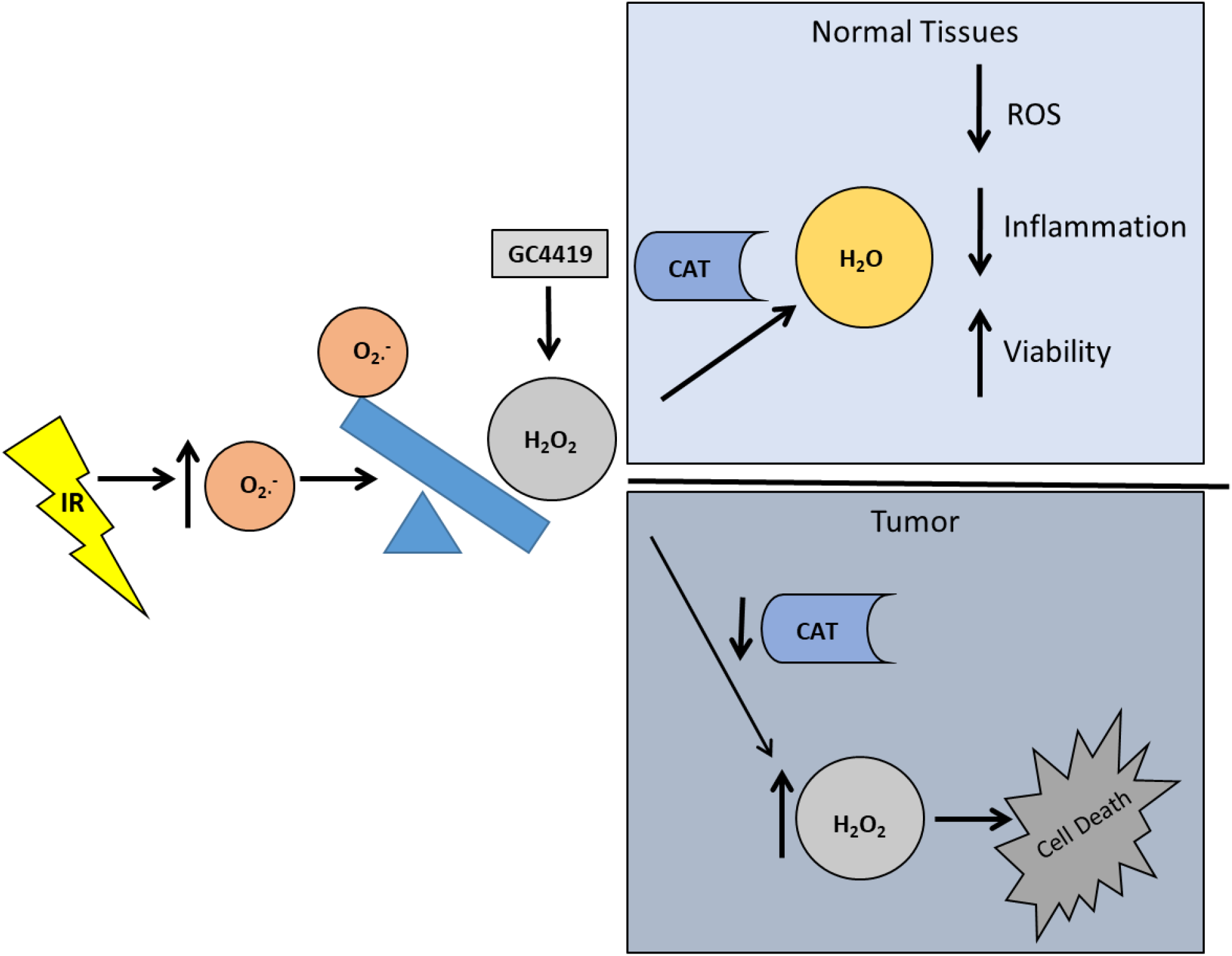
Model for dual efficacy of GC4419 as a radioprotector and enhancer of the radiation response. Ionizing radiation increases superoxide levels at the time of irradiation and upregulates biological superoxide production chronically. GC4419 dismutates superoxide so rapidly, that the concentration of hydrogen peroxide spikes. In normal tissue, high levels of catalase convert that hydrogen peroxide into water, reducing ROS and inflammation while increasing viability. In the tumor where catalase levels are repressed, hydrogen peroxide accumulates and produces cytotoxic effects that result in cell death.

